# Ventricular proximity associates with survival and immune remodelling in glioblastoma

**DOI:** 10.64898/2026.11.13.766477

**Authors:** Julia L. Gutiérrez-Arroyo, Chaitali Chakraborty, Pia Gallego-Porcar, Jaume Barceló, Clément Guillou, Pascal Cosset, Elvira Carbonell-Martínez, Marwen Ben Fdilen, Elena Tomás-Bort, Luis G. González-Bonet, Oscar Maiques, Silvia Remeseiro, Conrado Martínez-Cadenas, María Ángeles Marqués-Torrejón

## Abstract

Glioblastoma is clinically and molecularly heterogeneous, yet whether its position relative to the lateral ventricles shapes tumour biology remains unclear. Here we measured tumour–ventricle distance in 57 patients with IDH-wildtype glioblastoma and combined survival analysis with RNA sequencing, whole-genome sequencing, proteomics and multiplex immunofluorescence. Proximity to the ventricles was associated with shorter survival (log-rank p = 0.0012; univariable HR per 5-mm increase, 0.55; 95% CI 0.38–0.80). Ventricle-proximal tumours lacked a distinct genomic driver but displayed an inflammatory yet immunoregulatory profile: antigen-presentation, leukocyte-activation and cytokine programmes coexisted with checkpoint and immunosuppressive transcripts, and proteomics revealed concordant complement and myeloid signals. In tissue, these tumours tended to show more Ki67-positive cells and proliferating SOX2-positive stem-like cells, suggesting increased plasticity of the stem-like compartment, without more CD4- or CD8a-positive lymphocytes. Ventricular proximity thus marks a clinically adverse context with a distinct immune-inflammatory profile, providing a framework for validation and patient stratification.

## INTRODUCTION

Glioblastoma is an aggressive adult-type diffuse glioma, classified as IDH-wildtype, CNS WHO grade 4 under the 2021 World Health Organization criteria [1]. Despite surgery, radiotherapy and temozolomide-based treatment, survival is generally limited and recurrence is frequent [2]. Large-scale genomic studies have identified recurrent alterations involving receptor tyrosine kinase, PI3K, p53 and RB pathways, but these changes do not fully capture the variation in clinical behaviour among patients [3,4].

Glioblastoma also contains diverse malignant-cell states and a complex microenvironment. Immune cells, vascular and extracellular-matrix structures, hypoxia and resident neural populations contribute to heterogeneity within and between tumours [5–11]. This raises the possibility that anatomical location influences the tumour ecosystem alongside its genetic background.

The lateral ventricular region is of particular interest because of its proximity to subventricular neural progenitor niches, cerebrospinal-fluid interfaces and specialised tissue compartments [12–14]. Several studies associate ventricular contact with poorer survival [15,16], whereas a large two-cohort analysis found that, after adjustment for established prognostic factors, distance from the ventricle among non-contacting tumours did not correlate with survival [17]. Binary contact classifications may also overlook variation among tumours that do not directly about the ventricle. Measuring tumour–ventricle distance offers a quantitative way to investigate this spatial relationship.

The biological correlates of ventricular proximity remain uncertain. Differences could involve tumour-intrinsic variation, malignant-cell states or immune and stromal programmes. RNA sequencing, whole-genome sequencing and proteomics provide complementary molecular readouts, while multiplex immunofluorescence allows selected cellular markers to be assessed in preserved tissue architecture [18–20]. Because these platforms were applied to partially overlapping subcohorts, concordance across assays must be interpreted at the pathway level rather than as fully matched patient-level multi-omic validation.

We therefore examined whether tumour–ventricle distance was associated with survival in a clinical glioblastoma cohort and characterised related molecular features using RNA sequencing, whole-genome sequencing, label-free proteomics and tissue-based multiplex immunofluorescence. Our aim was not to define a new anatomical subtype, but to assess whether proximity to the ventricle accompanies reproducible patterns of tumour and microenvironmental biology.

## RESULTS

### Quantitative tumour–ventricle distance is associated with overall survival

The clinical cohort comprised 57 patients with IDH-wildtype glioblastoma (median age, 64 years; recorded range, 17–81 years), including 27 women and 30 men (Table 1). Minimum tumour–ventricle distance was measured on preoperative contrast-enhanced MRI between the enhancing tumour margin and the lateral ventricular ependymal surface (Fig. 1A–C). Cohort-derived distance groups were defined as Close (0–5.2 mm, n = 21), Intermediate (>5.2–12.5 mm, n = 17) and Far (>12.5 mm, n = 19).

**Table 1:** Characteristics of the clinical cohort.

| **Characteristic** | **Value** |
|---|---|
| IDH-wildtype glioblastoma | 57 patients |
| Age at diagnosis, median | 64 years |
| Female / male | 27 / 30 |
| Close / Intermediate / Far | 21 / 17 / 19 |
| Biopsy / partial / total resection | 18 / 26 / 13 |
| Progression/recurrence documented | 25 / 57 (43.9%) |

Values reflect the 57-patient clinical cohort.

**Figure 1:**
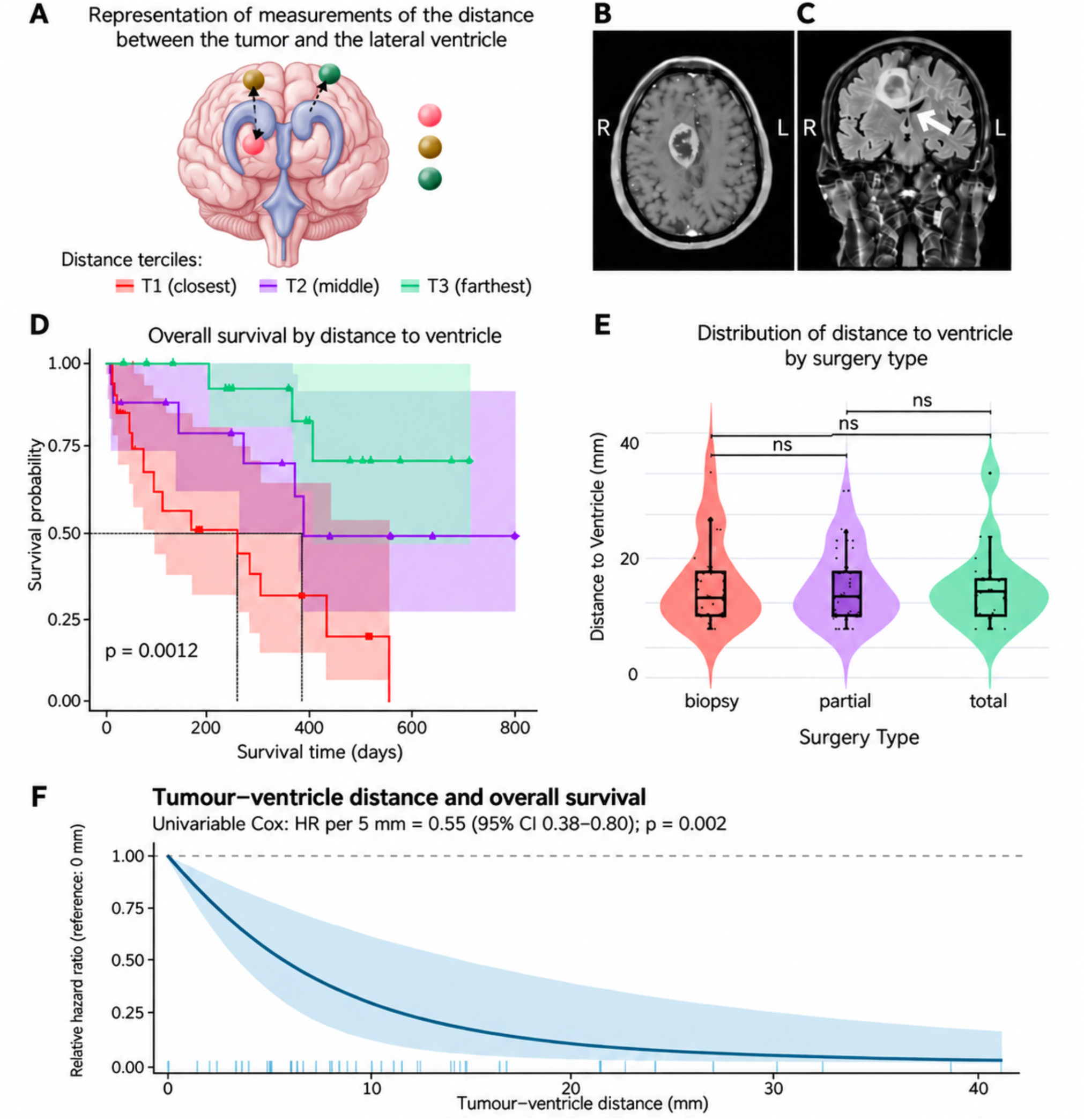
Tumour–ventricle distance and overall survival. (A) Schematic illustrating distance measurement in relation to the lateral ventricles and the three distance categories. (B,C) Representative axial and coronal magnetic resonance images indicating the tumour–ventricle measurement. (D) Kaplan–Meier overall-survival curves for Close (n = 21), Intermediate (n = 17) and Far (n = 19) tumours (log-rank p = 0.0012); shaded regions indicate 95% confidence intervals. (E) Distance distributions by surgical category (biopsy, partial resection and total resection; Kruskal–Wallis p = 0.91; pairwise Wilcoxon tests with Bonferroni correction showed no significant differences). (F) Univariable Cox analysis of distance on a continuous scale (HR per 5 mm = 0.55; 95% CI, 0.38–0.80; p = 0.002); the line depicts the fitted relative hazard (reference, 0 mm) with its 95% confidence interval; ticks mark individual tumour distances (n = 52 patients with complete data).

Overall survival differed across the three distance groups (log-rank p = 0.0012; Fig. 1D). The Close group showed the shortest survival, while median overall survival was not reached for the Far group within the observed follow-up. In a univariable Cox model, greater distance was associated with a lower hazard of death (HR per 5-mm increase, 0.55; 95% CI, 0.38–0.80; p = 0.002; Fig. 1F). The Cox model was fitted in the 52 patients with complete data (Methods).

Tumour volume showed no statistically significant association with overall survival when analysed by tertiles (p = 0.42), and the reported continuous-volume analysis also yielded a nonsignificant result (p = 0.85). Tumour volume and ventricular distance were not significantly associated (p = 0.53). Ventricular distance did not differ significantly across biopsy, partial resection and total resection categories (Kruskal–Wallis p = 0.91; Fig. 1E). Extent of resection showed a nonsignificant survival trend (log-rank p = 0.079), despite its established prognostic value in larger series [21]. These univariable findings support an association between ventricular distance and survival, but do not rule out confounding by other clinical variables.

### Ventricular proximity is not associated with a single global transcriptional subtype

Clinical and molecular data were available for 23 patients, of whom 22 passed RNA-seq quality control (Close, n = 6; Intermediate, n = 9; Far, n = 7). Overall survival differed significantly across ventricular-distance groups within this molecular subgroup (log-rank p = 0.0063). Despite this survival association, global transcriptomic analyses revealed substantial inter-patient heterogeneity, without a clear separation of tumours according to ventricular proximity. These findings suggest that the clinical differences associated with tumour location are not accompanied by a single, well-defined transcriptional subtype.

To investigate distance-associated transcriptional differences, we performed pairwise differential expression analyses with DESeq2 (Fig. 2A–F). At a Benjamini–Hochberg-adjusted p ≤ 0.01, we identified 819 differentially expressed genes (DEGs) between Close and Far tumours (299 upregulated and 520 downregulated in Close), 1,077 between Close and Intermediate tumours (651 upregulated and 426 downregulated in Close) and 468 between Intermediate and Far tumours (315 upregulated and 153 downregulated in Intermediate).

**Figure 2:**
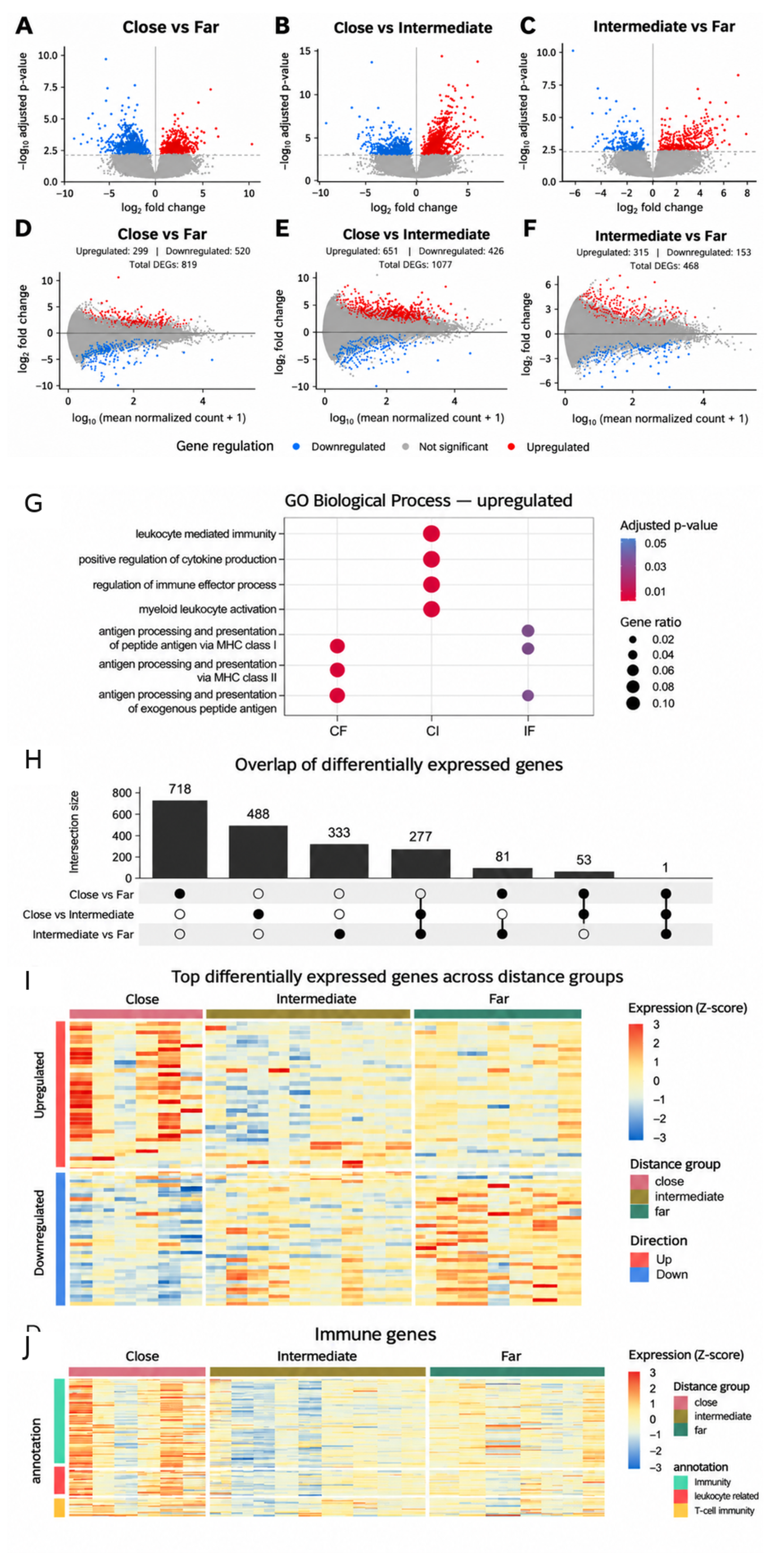
Differential gene expression and immune-associated transcriptional programmes according to ventricular proximity in glioblastoma. (A–C) Volcano plots showing differentially expressed genes (DEGs) in Close versus Far (A), Close versus Intermediate (B) and Intermediate versus Far (C). Red and blue dots represent upregulated and downregulated genes, respectively, whereas grey dots indicate non-significant genes. The horizontal dashed line indicates the adjusted significance threshold. (D–F) MA plots showing the relationship between mean normalised gene expression (log₁₀ scale) and log₂ fold change for the same comparisons. Red and blue dots indicate significantly upregulated and downregulated genes, respectively. The numbers of DEGs were 819 for Close versus Far (299 upregulated; 520 downregulated), 1,077 for Close versus Intermediate (651 upregulated; 426 downregulated) and 468 for Intermediate versus Far (315 upregulated; 153 downregulated). (G) Gene Ontology (GO) enrichment analysis of biological processes associated with upregulated genes across the three comparisons (CF, Close versus Far; CI, Close versus Intermediate; IF, Intermediate versus Far). Dot colour represents the adjusted p-value, and dot size indicates the gene ratio. (H) UpSet plot illustrating the intersections of DEGs identified across the three pairwise comparisons. Bar heights indicate intersection sizes, and connected filled circles identify the corresponding contrasts. (I) Heatmap of selected highly differentially expressed genes across Close, Intermediate and Far tumours, with rows classified as upregulated or downregulated. (J) Heatmap of immune-associated genes, including immune-response, leukocyte-related and T-cell immunity annotations. Heatmap colours represent row-scaled expression values (Z-scores). Differential expression was analysed using DESeq2, with statistical significance defined as a Benjamini–Hochberg-adjusted p-value ≤ 0.01.

The volcano plots (Fig. 2A–C) illustrate the magnitude and statistical significance of transcriptional changes across the three comparisons, highlighting distinct patterns of upregulated and downregulated genes. Consistently, the corresponding MA plots (Fig. 2D–F) demonstrate that differential expression occurs across a range of transcript abundance levels rather than being restricted to highly expressed genes. Together, these analyses indicate that ventricular-distance groups exhibit substantial transcriptional differences despite the absence of a distinct global expression profile.

To assess the overlap between transcriptional responses, an UpSet analysis was performed (Fig. 2H). Most DEGs were specific to individual pairwise comparisons (488 for Close versus Far, 718 for Close versus Intermediate and 333 for Intermediate versus Far). Smaller intersections were observed between pairs of comparisons (277 genes shared by Close versus Far and Close versus Intermediate, 81 by Close versus Intermediate and Intermediate versus Far, and 53 by Close versus Far and Intermediate versus Far), and only one gene was shared across all three contrasts. This limited overlap highlights the heterogeneity of transcriptional differences associated with ventricular distance.

The heatmap of selected DEGs further illustrated these patterns, revealing heterogeneous expression profiles across individual tumours and differences in the distribution of upregulated and downregulated genes among Close, Intermediate and Far samples (Fig. 2I). Overall, these findings support a relationship between ventricular proximity and specific transcriptional programmes rather than the existence of a uniform, distance-defined molecular subtype.

Close tumours exhibit immune-associated transcriptional activation alongside immunoregulatory programmes

To investigate the biological processes underlying these transcriptional differences, Gene Ontology (GO) enrichment analysis was performed on upregulated genes from each pairwise comparison (Fig. 2G). Immune-associated pathways were prominent, particularly in comparisons involving the Close group.

The Close versus Far comparison revealed enrichment of biological processes related to antigen processing and presentation, via both MHC class I and class II. In contrast, the Close versus Intermediate comparison was characterised by enrichment of leukocyte-mediated immunity, regulation of immune effector processes, myeloid leukocyte activation and positive regulation of cytokine production. Antigen-presentation-related processes were also represented in the Intermediate versus Far comparison. These findings indicate that ventricular-distance groups differ in immune-associated transcriptional programmes, with particularly prominent inflammatory and immune-regulatory signatures in comparisons involving tumours located closer to the ventricles.

The immune-associated heatmap (Fig. 2J) further revealed heterogeneous expression patterns among the three distance groups, including genes involved in general immune responses, leukocyte-related functions and T-cell immunity. Close tumours exhibited higher expression of several cytotoxicity- and immune-activation-associated transcripts, including CD8A, CD8B, GZMA, GZMB, PRF1 and IFNG (Fig. 2J). However, these activation-associated signals co-occurred with increased expression of transcripts implicated in inhibitory or immunoregulatory programmes, including LAG3, HAVCR2, TOX, ENTPD1, IDO1 and VSIR.

This coexistence of inflammatory and inhibitory transcriptional signatures suggests that enhanced immune-related activity in ventricular-proximal tumours may not necessarily translate into an effective antitumour response. Instead, it may reflect an immunologically active but potentially immunoregulatory tumour microenvironment. Nevertheless, bulk RNA-seq does not establish whether these transcripts originate from the same cell populations, nor can it directly demonstrate T-cell exhaustion or impaired immune function [22,23].

Taken together, the differential expression, functional enrichment and immune-associated transcriptional analyses suggest that ventricular proximity is associated with heterogeneous changes in immune and inflammatory programmes. The coexistence of cytotoxicity-related and immune-inhibitory transcripts raises the possibility that ventricular-proximal tumours harbour a microenvironment in which inflammatory activation occurs alongside mechanisms that may constrain effective antitumour immunity. These findings provide a molecular basis for further investigating the relationship between ventricular proximity, immune regulation and glioblastoma cellular plasticity.

### Deconvolution reveals heterogeneous malignant-cell states and immune composition

To further characterise the cellular composition and immune-associated transcriptional programmes of glioblastomas according to ventricular proximity, exploratory deconvolution analyses were performed using three complementary reference-based approaches (Fig. 3A–C).

**Figure 3:**
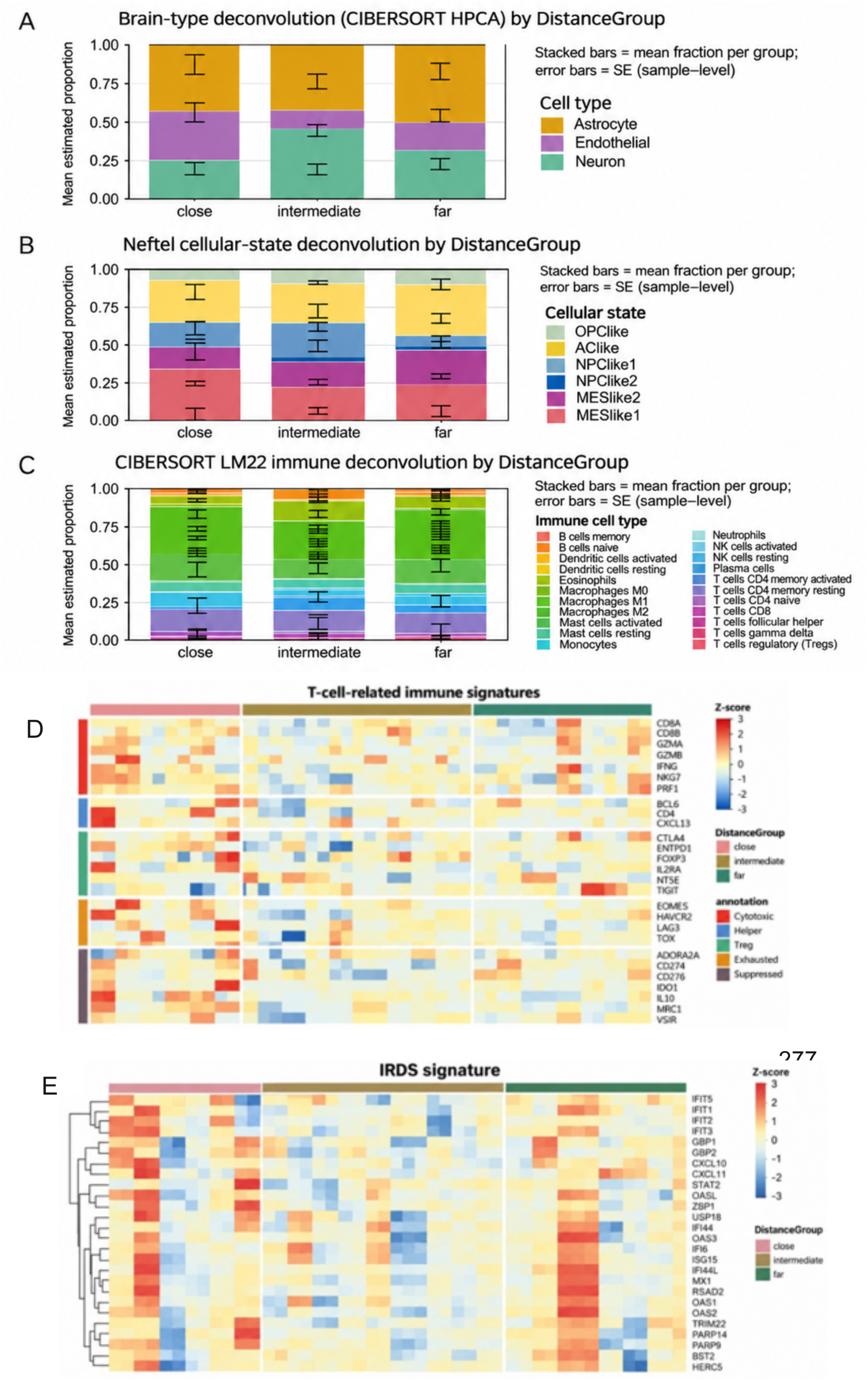
Exploratory deconvolution and immune-gene signatures derived from bulk RNA sequencing. (A) HPCA-based brain-associated cell-type estimates (neuron, endothelial and astrocyte). (B) Neftel-derived malignant-cell state estimates (MES-like, NPC-like, AC-like and OPC-like states). (C) CIBERSORT LM22 estimates for leukocyte populations. In A–C, stacked bars indicate group-mean estimated proportions for Close, Intermediate and Far; error bars denote standard errors (SE) calculated across biological samples, as indicated in the plots. (D) Heatmap of selected T-cell-related transcripts annotated by functional gene categories. (E) Heatmap of an interferon-related DNA-damage-resistance gene signature (IRDS). Heatmap colours indicate row-scaled expression (Z-scores); upper annotations identify distance groups. Deconvolution comparisons were exploratory and none remained significant after FDR correction.

HPCA-based deconvolution revealed differences in the estimated proportions of brain-associated cell populations across the three distance groups (Fig. 3A). Intermediate tumours exhibited a relatively higher estimated neuronal fraction, whereas Close tumours showed a greater contribution from endothelial-associated signatures compared with the other groups. Astrocyte-associated signatures were represented across all three categories, with considerable variation between patients. These findings suggest differences in brain-associated cellular programmes, although reference-based estimates from bulk RNA-seq cannot establish the anatomical origin or precise abundance of individual cell populations.

Deconvolution using Neftel-derived malignant-cell state signatures further highlighted transcriptional heterogeneity across ventricular-distance groups (Fig. 3B). Close tumours exhibited a higher mean estimated MES-like 1 fraction (33.77%) than Intermediate (21.67%) and Far (22.07%) tumours, whereas AC-like signatures were most prominent in the Far group (33.28%). NPC-like and OPC-like programmes were also detected across all groups, supporting the coexistence of multiple malignant transcriptional states rather than a single distance-associated phenotype. However, these differences did not remain statistically significant after false-discovery-rate correction.

Immune-cell deconvolution using CIBERSORT LM22 revealed heterogeneous leukocyte-associated profiles across the three groups (Fig. 3C). Macrophage-related signatures, including M0, M1 and M2 populations, represented a substantial component of the estimated immune-cell composition, while additional signatures associated with NK cells, T lymphocytes, monocytes and other leukocyte populations were detected at varying proportions. The presence of M2-associated signatures is compatible with immunoregulatory myeloid programmes; nevertheless, these computational estimates cannot establish macrophage polarisation or functional immunosuppression. No individual immune-cell population consistently distinguished the ventricular-distance groups following multiple-testing correction.

To investigate whether these cellular estimates were accompanied by differences in immune-associated gene expression, we examined selected T-cell-related transcriptional signatures (Fig. 3D). The heatmap revealed heterogeneous expression of genes associated with cytotoxic activity (CD8A, CD8B, GZMB, GZMA, IFNG and PRF1), helper and regulatory functions, and inhibitory or exhaustion-associated programmes, including LAG3, HAVCR2, TOX, ENTPD1, IDO1 and VSIR. Genes upregulated in Close tumours in the Close-versus-Far and Close-versus-Intermediate comparisons also included HLA-DRA, IL10, CD28, IL2RA, ICOS, CXCL10 and CXCR3. Several Close tumours exhibited increased expression of cytotoxicity-associated transcripts alongside inhibitory and immunoregulatory markers, suggesting that immune activation may coexist with mechanisms potentially limiting effective antitumour responses. However, the expression patterns varied considerably between patients, and bulk transcriptomic data cannot determine whether these genes are co-expressed within individual immune cells or demonstrate functional T-cell exhaustion.

Finally, we evaluated the interferon-related DNA-damage-resistance signature (IRDS), which comprises genes involved in interferon signalling and stress-associated cellular responses (Fig. 3E). Several interferon-stimulated genes, including IFIT family members, ISG15, MX1, OAS family members and STAT1, displayed increased expression in subsets of Close tumours. Nevertheless, the signature was heterogeneous, with elevated expression also observed in individual Far tumours and lower expression in other Close samples. These observations suggest that interferon-associated transcriptional programmes may be particularly prominent in certain ventricular-proximal glioblastomas but do not represent a uniform characteristic of the Close group. Furthermore, enrichment of an IRDS-associated transcriptional signature should not be interpreted as direct evidence of treatment resistance without functional validation.

Collectively, the deconvolution and immune-signature analyses reveal heterogeneous brain-associated, malignant-cell and immune-related transcriptional programmes across ventricular-distance groups. The coexistence of inflammatory, myeloid-associated, mesenchymal-like and immunoregulatory signatures in subsets of ventricular-proximal tumours supports further investigation of the relationship between anatomical location, immune regulation and malignant cellular plasticity. However, these exploratory findings do not establish a distinct cellular composition or functional immune state specific to periventricular glioblastoma.

### Whole-genome sequencing does not define a periventricular genomic driver

Whole-genome sequencing was performed on tumour and matched blood DNA. Quality-control metrics were available for 22 DNA samples, with reported median unique alignment of 94.5% and a median per-sample mean coverage of 59.3× (range, 54.6–145.1×). The genomic-region distributions of somatic single-nucleotide variants (SNVs) and indels varied across ventricular-distance groups (Fig. 4A,B), but no recurrent somatic alteration uniquely defined the Close group. The distributional contrasts were therefore considered descriptive and exploratory rather than evidence for a distinct genomic subtype.

**Figure 4:**
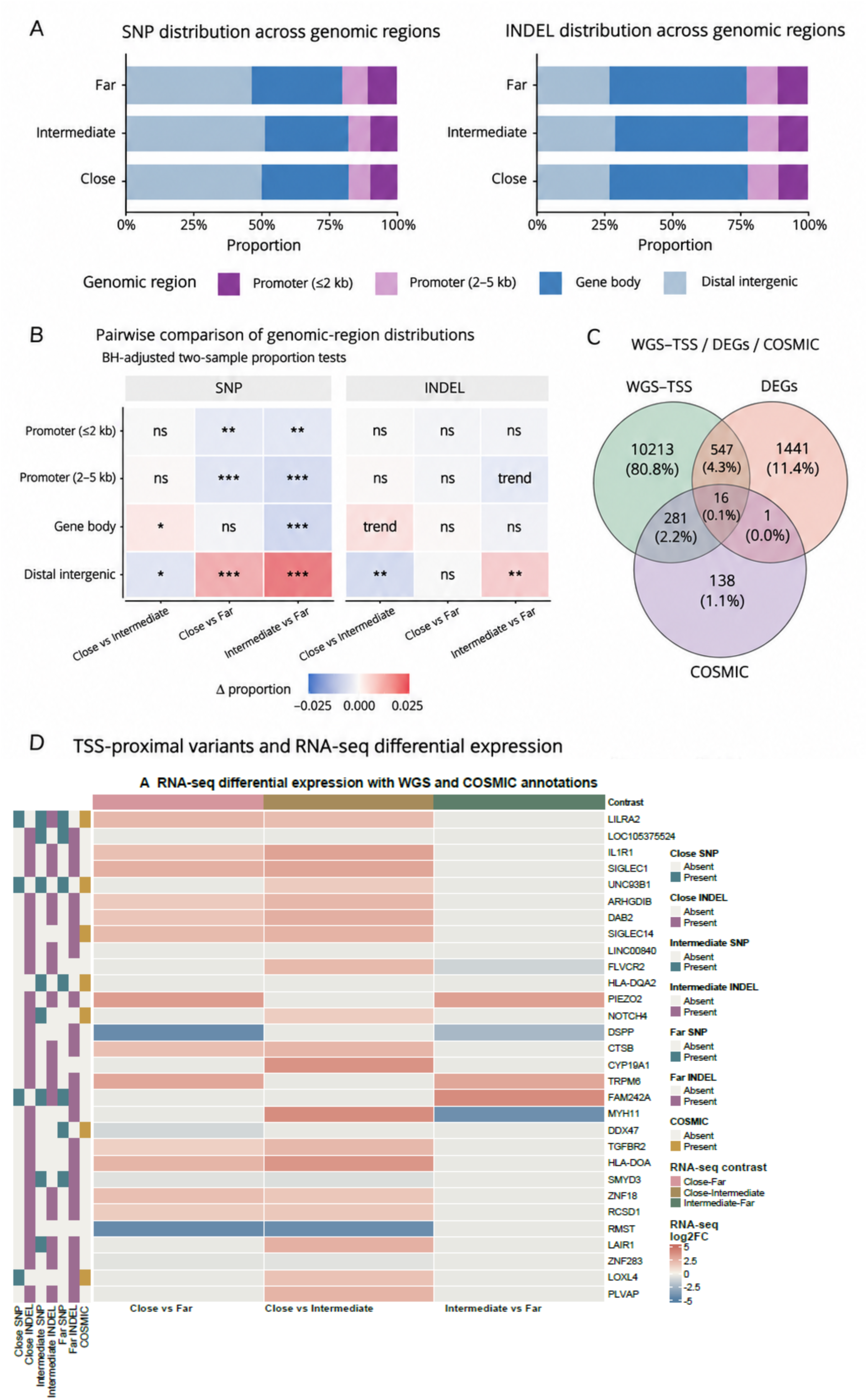
WGS variant distributions and integration with RNA-seq. (A) Relative proportions of SNPs (somatic SNVs) and INDELs assigned to distal intergenic regions, gene bodies and promoter regions (≤2 kb and 2–5 kb from the TSS) in Close, Intermediate and Far tumours. (B) Pairwise comparisons of region-specific variant proportions for SNPs and INDELs; tile colours represent differences in proportions and asterisks denote BH-adjusted significance of two-sample proportion tests (*p < 0.05, **p < 0.01, ***p < 0.001; trend, p < 0.10). (C) Three-set Venn diagram showing overlap among genes associated with TSS-proximal WGS variants, DEGs and COSMIC-annotated genes; numbers give set intersections. (D) Heatmap of RNA-seq log2 fold changes for selected genes across the three contrasts, with annotation tracks indicating SNP, INDEL and COSMIC overlap. All TSS-proximal associations are candidate-level and do not imply a direct regulatory effect.

To explore possible regulatory candidates, variants within ±10 kb of annotated transcription start sites (TSSs) were intersected with RNA-seq DEGs and COSMIC annotations (Fig. 4C,D). Candidate genes included LILRA2, IL1R1, SIGLEC1, ARHGDIB, DAB2, SIGLEC14, LINC00840 and FLVCR2. Although several are implicated in innate immunity or myeloid-associated processes, overlap between a nearby variant and differential expression does not demonstrate that the variant altered transcription.

Across the candidate lists, immune-related and myeloid-associated genes recurred in different contrasts, but their presence was not uniform across samples. Together, the WGS and RNA-seq data suggest candidates for further investigation, without establishing a distance-specific genomic driver or a causal regulatory mechanism [3,4].

### Proteomics highlights inflammatory and myeloid-associated processes in Close tumours

The proteomic cohort included nine initially processed tumours; eight independent samples passed quality control (Close n = 3, Intermediate n = 2, Far n = 3). After requiring at least two identified peptides per protein, 1,505 proteins were retained. An initial Progenesis analysis, in which technical replicates were treated as separate measurements, flagged 378 proteins as differentially abundant in at least one comparison; PCA of this selected protein set showed separation of many Close and Far samples along PC1 (47.3% of the variance), whereas the Intermediate samples were heterogeneous (Fig. 5A,B). This unsupervised display of a preselected protein set should not be interpreted as independent validation of group separation.

**Figure 5:**
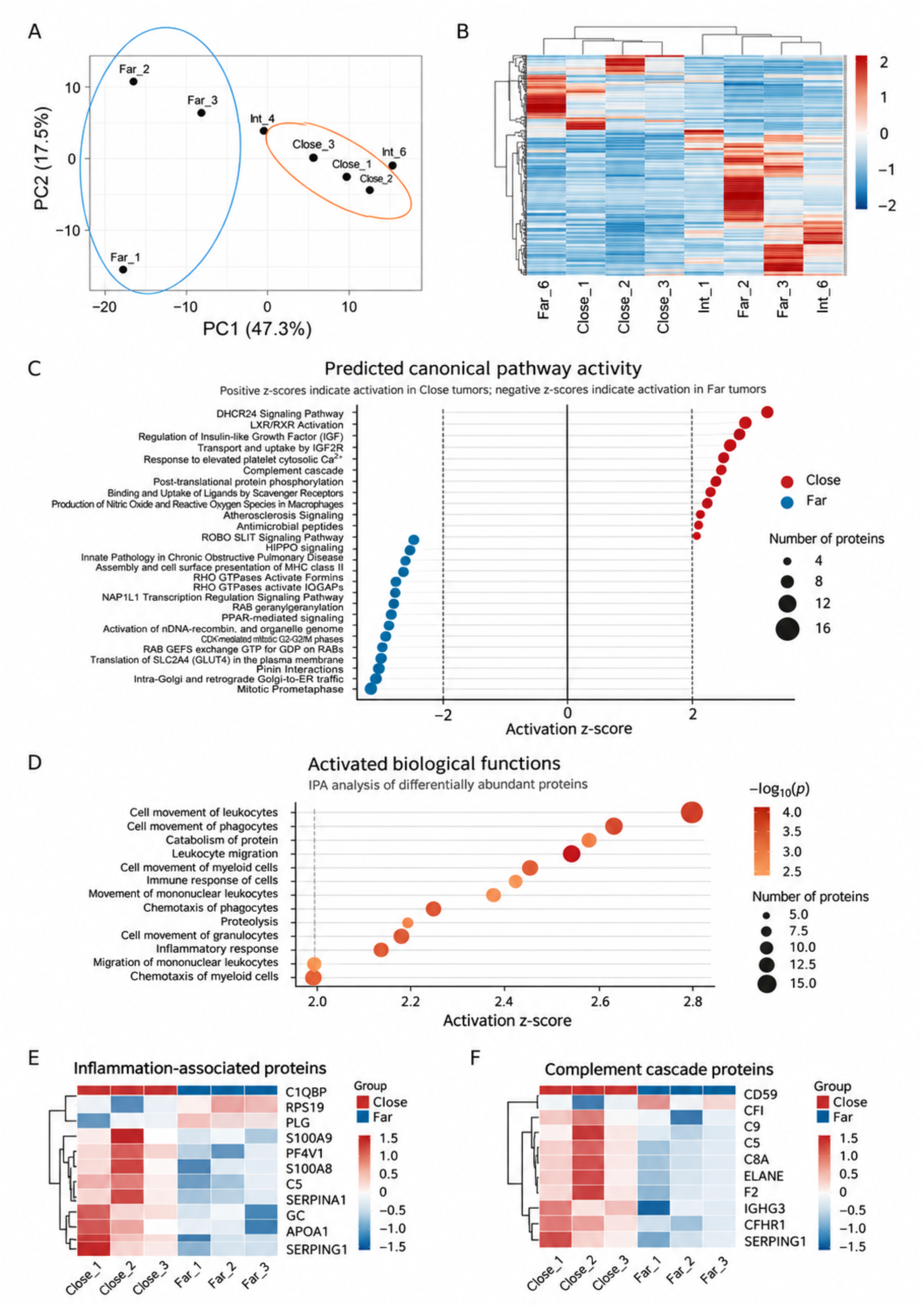
Exploratory proteomic profiles according to ventricular distance. (A) Principal-component analysis of the selected protein-abundance dataset; PC1 and PC2 explain 47.3% and 17.5% of the plotted variation, respectively. (B) Unsupervised hierarchical heatmap of protein abundances across the eight QC-passing tumour specimens (Close n = 3, Intermediate n = 2, Far n = 3). (C) IPA-predicted canonical pathway activity, with positive and negative Z-scores representing the predicted directional contrast as labelled in the panel; point size indicates the number of contributing proteins. (D) IPA-predicted biological functions, shown by activation Z-score, nominal significance and number of associated proteins. (E) Heatmap of selected inflammation-associated proteins and (F) complement-related proteins in Close and Far samples. These exploratory analyses are based on a small number of independent tumours; IPA Z-scores denote predictions, not direct functional assays.

The exploratory Close-versus-Far comparison identified 193 proteins meeting nominal p < 0.05 and the specified fold-change threshold (|log2FC| ≥ 1), of which 67 were more abundant in Close and 126 in Far tumours. Proteins with higher abundance in Close tumours included S100A8, S100A9, C5, CFHR1, SERPING1, ELANE and SERPINA1; RPS19, C1QBP and CD59 were relatively enriched in Far tumours. Given the three independent samples per group, these findings should be interpreted as candidate-level observations.

Ingenuity Pathway Analysis (IPA) suggested increased activity of pathways and functions related to inflammatory responses, leukocyte movement, phagocyte migration, complement and myeloid biology, including predicted macrophage nitric oxide and reactive-oxygen-species production, in the Close–Far comparison (Fig. 5C,D). Inflammation-associated and complement-related protein subsets are illustrated in the corresponding heatmaps (Fig. 5E,F). IPA activation Z-scores are computational predictions based on protein-abundance patterns, not direct measurements of pathway activation or macrophage function.

### Multiplex immunofluorescence shows heterogeneous tissue-level marker profiles

Multiplex immunofluorescence was analysed in 12 FFPE glioblastoma sections (four tumours per group; Fig. 6). Cell DIVE-derived outputs included the fraction of segmented cells positive for SOX2, CD163, CD68, SMA, COL1A1, Ki67, CD4 and CD8a, as well as proliferative and non-proliferative SOX2-positive subsets. The number of segmented cells ranged from 23,344 to 1,624,536 across specimens. As each tissue, rather than each segmented cell, is the independent biological sample, marker percentages were summarised at the specimen level and displayed as individual points with group means and standard deviations (Fig. 6A).

**Figure 6:**
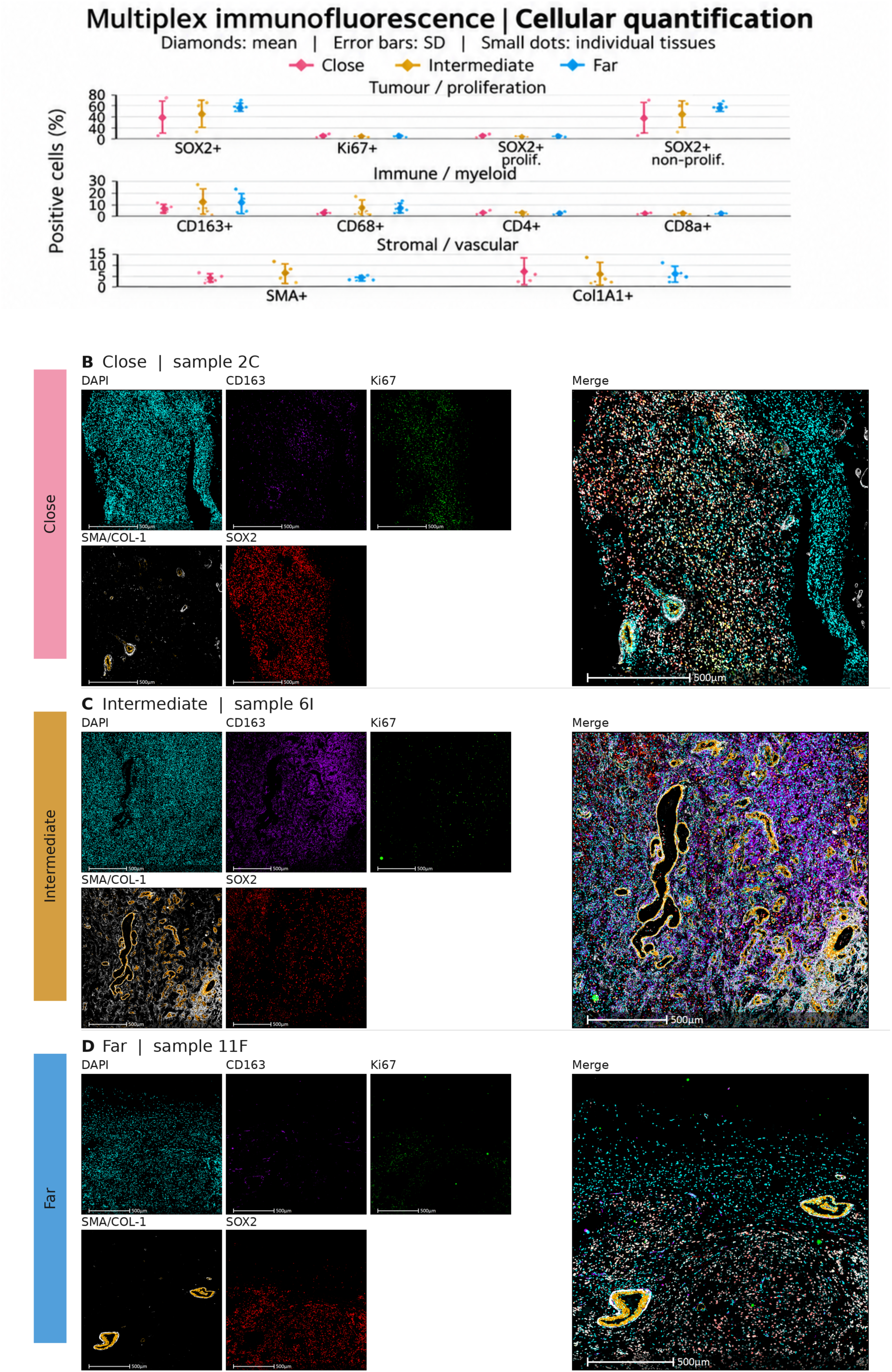
Multiplex immunofluorescence reveals heterogeneous cellular marker profiles across ventricular-distance groups. (A) Quantification of marker-positive cells in Close, Intermediate and Far glioblastomas, grouped into tumour-associated and proliferative markers (SOX2, Ki67, proliferative SOX2-positive and non-proliferative SOX2-positive cells), immune and myeloid markers (CD163, CD68, CD4 and CD8a), and stromal and vascular markers (SMA and COL1A1). Small dots represent individual tumour samples (n = 4 biological specimens per group), diamonds indicate group means, and error bars represent ± standard deviation (SD), with lower and upper limits truncated to the biologically valid range of 0–100% where applicable. Groups are shown in pink (Close), ochre (Intermediate) and blue (Far). (B–D) Representative multiplex immunofluorescence images from Close (B, sample 2C), Intermediate (C, sample 6I) and Far (D, sample 11F) tumours. Each montage displays DAPI (cyan), CD163 (purple), Ki67 (green), SMA/COL1 (white/yellow), SOX2 (red) and the merged fluorescence image. Scale bars, 500 µm. Quantitative measurements are presented at the biological-sample level, whereas representative images illustrate tissue-level spatial heterogeneity. Comparisons are descriptive; no inferential tests were performed.

The mean SOX2-positive fraction was 38.62 ± 28.48% in Close, 45.13 ± 23.23% in Intermediate and 58.29 ± 5.52% in Far tumours. Most SOX2-positive cells were classified as non-proliferative based on the absence of Ki67 staining at the time of measurement; this classification does not establish a quiescent, reversible GSC state. In contrast to total SOX2 positivity, the mean proliferative SOX2-positive fraction was highest in Close tumours (2.58 ± 2.88%), compared with Intermediate (0.59 ± 0.48%) and Far (1.58 ± 1.93%). Ki67 positivity showed a comparable descriptive pattern (3.31 ± 3.10%, 1.12 ± 0.51% and 1.97 ± 2.18%, respectively). These findings describe variation in marker-positive fractions, without statistically established differences between groups or direct evidence of transitions between cell-cycle states.

Myeloid-associated marker fractions were variable and, on average, lower in Close tumours: CD163 positivity averaged 5.22%, 10.63% and 9.74% in Close, Intermediate and Far samples, respectively, while CD68 positivity averaged 1.57%, 5.13% and 5.10%. CD4 positivity was low in all groups (1.54%, 1.30% and 1.21%), as was CD8a positivity (0.82%, 0.69% and 0.82%). SMA and COL1A1 were also heterogeneous across samples (Fig. 6A). The representative microscopy fields show DAPI-labelled nuclei, CD163-positive signal, Ki67-positive nuclei, a combined SMA/COL1 channel, SOX2 staining and a merged composite (Fig. 6B–D). In this small series, the stronger immune- and myeloid-related RNA-seq and proteomic signatures near the ventricle did not correspond to higher lymphocyte- or macrophage-marker fractions.

## DISCUSSION

Earlier studies have linked lateral ventricular contact to poorer outcome in glioblastoma, although findings vary across cohorts and measurement approaches [15–17]. In the largest distance-based analysis, survival depended on ventricular contact rather than on distance among non-contacting tumours [17]; because our Close group (0–5.2 mm) includes contacting tumours, the graded association reported here may partly reflect that subset. We previously reported an association between tumour proximity to the ventricular system and survival in this patient cohort [24]. The contribution of the present study is therefore the analysis of distance as a continuous variable and the multi-omic characterisation of that cohort, not the prognostic association itself, and it is not an independent validation. The effect estimate derives from a univariable Cox model, which does not establish independence from age, treatment, resection extent or molecular characteristics; an adjusted analysis and external validation would be required before distance could be considered a prognostic classifier.

The biological relevance of the ventricular region may extend beyond simple anatomical contact. The subventricular zone provides a specialised neural progenitor environment, and neural stem cells have been proposed as one possible cell of origin for subsets of glioblastoma [12–14,25]. These concepts sit within a broader literature highlighting glioma stem cells, epigenetic plasticity and interactions with the tumour microenvironment as determinants of growth and treatment resistance [26]. Nevertheless, the proximity of a tumour to the lateral ventricle does not prove that it originated from the subventricular niche, nor does it establish direct exposure to specific niche-derived signals.

Our transcriptional analyses offer a possible biological context for the survival association. Although Close tumours did not form a discrete global expression cluster, selected comparisons were enriched for antigen-presentation, interferon-related, leukocyte-activation and cytokine-associated programmes, and cytotoxicity-associated transcripts appeared alongside immune-inhibitory and immunoregulatory genes. Such coexistence is biologically plausible, because inflammatory signalling can persist without producing an effective antitumour response [8,22,23], and it is not inconsistent with the limited efficacy of PD-1 blockade in unselected patients with recurrent glioblastoma [27]. Transcript abundance, however, cannot establish immune dysfunction, and the absence of paired functional assays, uncertainty about the cellular source of individual transcripts and marked inter-patient variability preclude describing the Close group as immunologically inactive or uniformly T-cell exhausted.

HLA class II genes were among the recurrent immune-related signals, together with LAG3 and IL10 among the genes upregulated in Close tumours. In human glioblastoma, high HLA class II expression has been associated with LAG-3-positive CD4 T-cell infiltration, increased IL-10 and poorer outcome [28]. Our data contain several components of this axis but do not demonstrate the interaction, and MHC class II-dependent antigen presentation is not inherently suppressive; its meaning depends on the surrounding immune context.

The proteomic data point in a similar direction. Proteins more abundant in Close tumours included S100A8, S100A9, C5, CFHR1, SERPING1, ELANE and SERPINA1. S100A8 and S100A9 are highly expressed by myeloid-derived suppressor cells in glioma and have been linked to arginase expression and T-cell suppression [29,30], and complement activity can support tumour-promoting myeloid phenotypes [31]. IPA additionally predicted increased macrophage nitric oxide and reactive-oxygen-species production in Close tumours. These are computational predictions: neither molecule was measured, and C5a–C5aR1 signalling was not assessed. Together, these observations are consistent with inflammatory myeloid activity that may coexist with restrained antitumour immunity, but they generate hypotheses rather than evidence of myeloid-mediated immunosuppression.

Estimated cell composition and malignant-cell states were heterogeneous across groups. The mean MES-like 1 fraction was higher in Close tumours, but these differences did not remain significant after multiple-testing correction. The exploratory deconvolution therefore suggests possible shifts in cell-state programmes rather than a reproducible cell-type enrichment. Importantly, mechanistic work has demonstrated that interactions with immune cells can promote mesenchymal-like transitions in glioblastoma, including macrophage-derived oncostatin M signalling through OSMR/LIFR–GP130–STAT3 [32], and mesenchymal differentiation has been linked to NF-κB activity and radiation resistance [33]. Our co-occurrence of myeloid-associated and MES-like signals is compatible with this framework, but we did not directly measure oncostatin M signalling or demonstrate a macrophage-driven transition in these specimens.

Whole-genome sequencing did not identify a single recurrent alteration characteristic of ventricular-proximal tumours. Intersections between TSS-proximal variants, RNA-seq DEGs and COSMIC annotations highlighted candidate genes associated with immune and myeloid pathways, but spatial coincidence between variants and genes does not establish transcriptional regulation, and linking variant carrier status to expression in matched samples would be required to test it. The partially unmatched samples limited biological replication and pathway-inference methods restrict the cross-platform observations to convergence at the level of biological themes [19].

Multiplex immunofluorescence adds an exploratory tissue-level perspective. CD4 and CD8a fractions were low across groups, and CD163 and CD68 fractions were heterogeneous and, on average, lower in Close tumours than in the other groups. This does not contradict the RNA-seq and proteomic signals: bulk approaches measure pathway activity across mixed cell populations, whereas multiplex imaging quantifies a restricted set of markers in sampled tissue regions. The data therefore favour differences in immune state over a simple increase in immune-cell abundance. Total SOX2 positivity was descriptively highest in Far tumours, whereas the fractions of Ki67-positive and proliferative SOX2-positive cells were highest on average in Close tumours, which distinguishes total SOX2 expression from proliferation within the SOX2-positive compartment. Neither SOX2 positivity nor Ki67 negativity is sufficient to establish a tumour-initiating or reversibly quiescent glioma stem-cell population.

The proliferative SOX2-positive pattern raises a testable question about stem-like cell-state transitions. FOXG1 and SOX2 have been shown to regulate neural stem-cell identity and cell-cycle programmes in glioblastoma [34], and FOXG1 cooperates with Wnt/β-catenin signalling to promote exit from quiescence in experimental stem-cell models [35]. Here, neither FOXG1 signaling nor Wnt activity were assessed, and static measurements cannot show that SOX2-positive, Ki67-negative cells reenter the cell cycle. An important direction for future work is to test whether the periventricular environment promotes such transitions, for example through lineage tracing or functional assays in patient-derived stem-like cells.

Integrating the clinical, transcriptomic, proteomic and tissue findings, we propose a working model in which proximity to the ventricular niche may be associated with an inflammatory yet potentially immunoregulatory ecosystem that permits malignant-cell plasticity. Antigen-presentation and checkpoint-associated signals, interferon and complement activity, and myeloid-associated programmes could coexist with ineffective tumour clearance and, potentially, microenvironmental cues favouring mesenchymal adaptation and shifts in stem-like cell state [22,23,32,34,35]. These processes might contribute to tumour persistence and offer one hypothesis for the poorer survival observed near the ventricle. Our data are observational, do not resolve causal direction and do not establish a mechanistic link between immune dysfunction and reactivation of quiescent stem-like cells; functional immune profiling, lineage tracing, spatially resolved analyses and experimental interrogation of the proposed pathways will be required to test the model.

If validated, tumour–ventricle distance is a readily measurable variable from routine imaging that could be explored as a stratification factor in prospective biomarker studies, including immunotherapy trials. The coexistence of checkpoint, antigen-presentation, complement and myeloid signals argues against a single therapeutic target in Close tumours; strategies addressing both lymphocyte dysfunction and myeloid-mediated immunosuppression may merit investigation in this subgroup [27]. Our findings do not support treating periventricular glioblastomas differently from other glioblastomas, and the present study cannot predict response to any specific treatment.

Several limitations constrain interpretation. The 57-patient clinical cohort and smaller molecular subcohorts were selected retrospectively, were not fully matched across assays and were not tested in an independent cohort; the clinical cohort has been reported previously [24]. The Cox model was univariable and used a smaller available-case set than the clinical cohort; residual confounding remains unresolved, and categorising distance with cohort-derived cut-offs reduces information and makes results dependent on those cut-offs. Distance was measured in two dimensions from the enhancing tumour margin rather than as a three-dimensional minimum, and sampled tissue may not represent the region nearest the ventricle. The transcriptomic data reflect bulk tissue and cannot assign expression to specific malignant or immune cells. Deconvolution is inferential. The proteomic series is small, and its initial feature selection treated technical replicates as separate measurements, whereas the multiplex series (four tumours per group) provides limited power for group comparisons. Ki67 and SOX2 staining cannot directly measure quiescence, immune-cell function or cell-state reactivation. Finally, tumour purity and immune/stromal components were assessed computationally from bulk RNA-seq, but these inferred measures were not independently validated by histological assessment. Variation in non-neoplastic tissue content between sampled regions could therefore contribute to some of the transcriptomic and proteomic patterns observed.

In conclusion, tumour proximity to the lateral ventricles was associated with poorer overall survival and distinct immune- and inflammation-related molecular programmes in IDH-wildtype glioblastoma. Rather than identifying a unique periventricular molecular subtype, our multi-omic analyses revealed substantial heterogeneity alongside convergent immune and myeloid-associated signals. Multiplex immunofluorescence further highlighted differences in proliferation-associated marker patterns, suggesting that ventricular proximity may be linked to changes in the tumour microenvironment and malignant-cell states. These findings provide a basis for future patient-matched, spatially resolved and functional studies aimed at understnding how anatomical location relates to glioblastoma progression (Figure 7).

**Figure 7:**
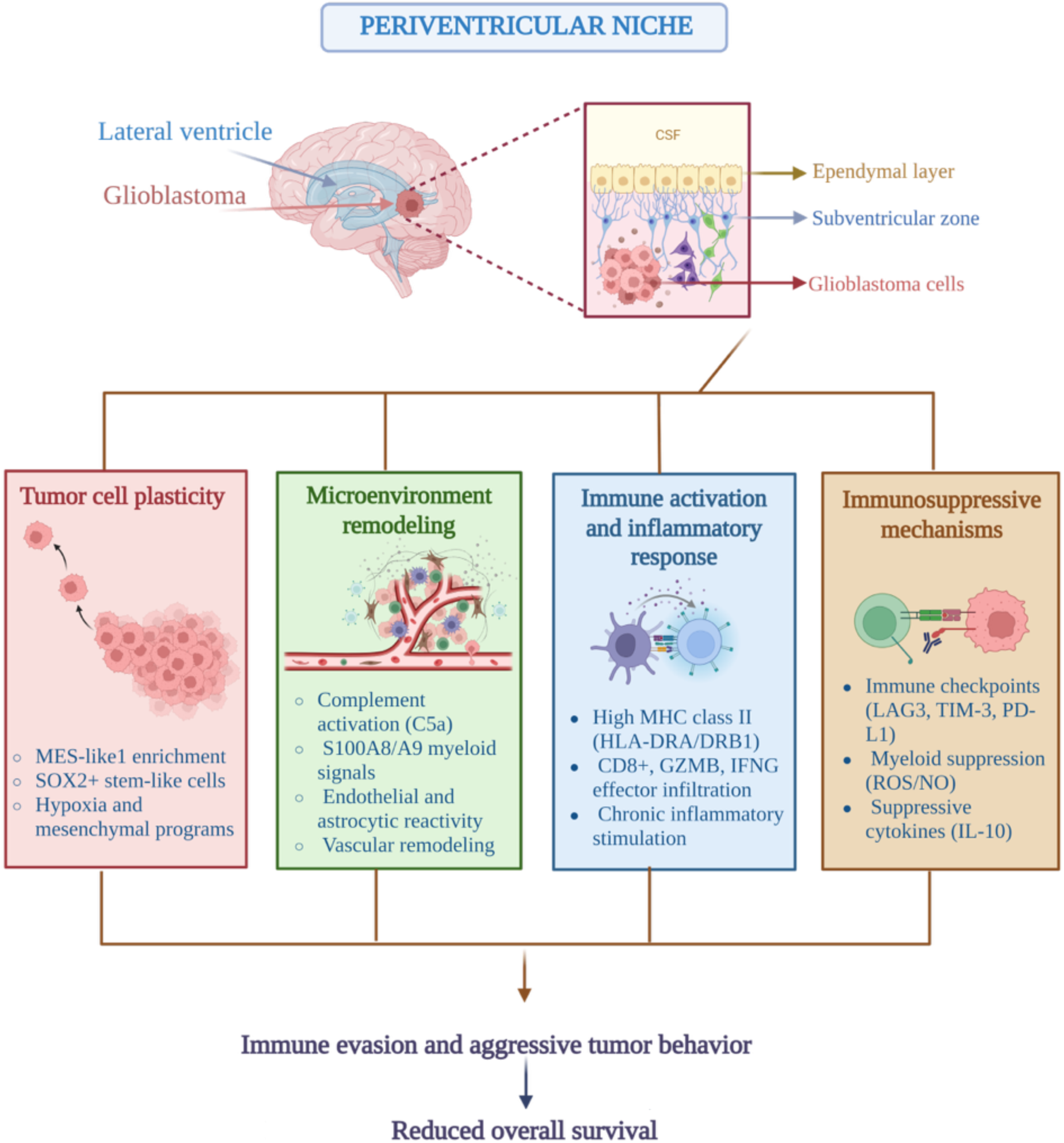
Proposed biological mechanisms associated with the periventricular glioblastoma microenvironment. Schematic representation of the lateral ventricular niche, including the ependymal layer, subventricular zone and adjacent glioblastoma cells. Multi-omic analyses suggest that ventricular proximity may be associated with four interconnected biological processes: tumour-cell plasticity, microenvironmental remodelling, immune and inflammatory activation, and immunoregulatory mechanisms. These include mesenchymal-like transcriptional programmes, myeloid- and complement-associated signalling, antigen presentation, cytotoxic immune signatures and immune checkpoint-related pathways. Together, these processes may contribute to a tumour microenvironment associated with aggressive glioblastoma behaviour and poorer overall survival. The proposed interactions are hypothesis-generating and do not establish direct causal mechanisms.

## METHODS

### Study cohort and ethics

This retrospective observational study included 57 patients with IDH-wildtype glioblastoma treated at the General University Hospital of Castellón, Spain. Classification followed the 2021 WHO criteria, and tumours with IDH1 or IDH2 mutations were excluded. Clinical and radiological information was derived from medical records. The study was approved by the Drug Research Ethics Committee (CEIm) of the General University Hospital of Castellón and conducted in accordance with the Declaration of Helsinki. Written informed consent for the use of tumour tissue and clinical data was obtained from participants. The study was conducted within the research project “Cazafantasmas-GBM” (protocol version 1, 17 January 2022), which received a favourable opinion from the Research Ethics Committee for Medicinal Products (CEIm) of the General University Hospital of Castellón (meeting of 31 January 2022, minutes 1/2022; certificate issued 9 March 2022). The study was conducted in accordance with the Declaration of Helsinki, and written informed consent for the use of tumour tissue and clinical data was obtained from all patients or their legal representatives. This cohort has been described in a previous clinical publication [24].

### Tumour–ventricle distance and clinical variables

Tumour–ventricle distance was measured on a selected preoperative contrast-enhanced MRI coronal section showing maximum tumour extension. The shortest linear distance from the enhancing tumour border to the ependymal surface of the lateral ventricle was recorded. This two-dimensional measure does not necessarily equal the true three-dimensional minimum distance and excludes non-enhancing infiltration. Analyses used distance as a continuous variable and as cohort-derived categories: Close (0–5.2 mm), Intermediate (>5.2–12.5 mm) and Far (>12.5 mm). Extent of resection was classified as biopsy, partial/subtotal or total resection. Overall survival was calculated from surgery to death or last follow-up.

### Sample selection and nucleic-acid extraction

Fresh-frozen tumour material was available for selected patients. RNA and DNA were isolated from approximately 20–30 mg tissue using the AllPrep DNA/RNA Mini Kit (QIAGEN). Concentration and purity were assessed using NanoDrop and Qubit assays; RNA integrity was evaluated using an Agilent Bioanalyzer. Platform-specific quality-control criteria were applied before RNA sequencing and WGS. The resulting analysis sets comprised 22 RNA-seq specimens and 22 DNA specimens with WGS quality-control metrics (Table 2).

**Table 2:** Analytical cohorts and limitations.

| **Platform** | **Biological samples** | **Interpretation** |
|---|---|---|
| Clinical | n = 57 | Survival analysis: Kaplan–Meier and univariable Cox (available cases n = 52) |
| RNA-seq | n = 22 (6/9/7) | DESeq2, Gene Ontology enrichment and exploratory deconvolution |
| WGS | n = 22 QC DNA samples | Somatic variants and TSS-proximal candidate integration |
| Proteomics | n = 8 (3/2/3) | Label-free profiling; exploratory contrasts and IPA predictions |
| Multiplex MxIF | n = 12 (4/4/4) | Cell-level positivity summarised at tissue level; descriptive only |

Clinical, molecular and imaging samples overlap only partially. Technical replicates were not counted as independent patients.

### RNA sequencing and transcriptomic analysis

Stranded, paired-end RNA sequencing (2 × 151 bp) was performed at the Centro Nacional de Análisis Genómico (Barcelona, Spain). Technical lane/index FASTQ files belonging to the same biological specimen were combined before alignment. Read quality was assessed using FastQC v0.12.1, and reads were aligned to GRCh38 using STAR v2.7.11b, with gene counts from STAR GeneCounts.

Differential expression was analysed in R with DESeq2 using proximity category as the grouping variable. Pairwise contrasts compared Close, Intermediate and Far groups, and genes with BH-adjusted p ≤ 0.01 were retained as DEGs. Gene Ontology analyses used clusterProfiler. Neftel-derived cellular-state signatures, an HPCA-derived brain-cell reference and CIBERSORT LM22 leukocyte signatures were applied as exploratory bulk RNA-seq deconvolution methods. Cell-state and cell-type estimates were summarised by patient, with Kruskal–Wallis tests and BH correction used for group-level exploratory comparisons.

### Whole-genome sequencing and genomic–transcriptomic integration

Whole-genome sequencing of tumour and matched peripheral-blood DNA was carried out at the Centro Nacional de Análisis Genómico. Reads were aligned to GRCh38 with BWA-MEM; duplicates were marked with Picard and base-quality-score recalibration used GATK tools. Somatic short variants were identified using Mutect2 and Strelka2 and annotated with VEP and ANNOVAR, with reference databases including gnomAD, ClinVar and COSMIC.

For genomic–transcriptomic integration, single-nucleotide variants (SNVs) and indels within ±10 kb of annotated transcription start sites were assigned to candidate genes and intersected with RNA-seq DEGs. COSMIC annotations were considered as an additional candidate-prioritisation layer. These overlaps identify candidates but cannot by themselves determine the direction or mechanism of regulation.

### Proteomic analysis

Fresh-frozen glioblastoma tissue from nine patients was initially processed for label-free proteomics; eight biological samples were retained after quality control (three Close, two Intermediate and three Far). Frozen tissue was mechanically disrupted in denaturing lysis buffer containing 7 M urea, 2 M thiourea, 20 mM DTT, 2% CHAPS and 0.5% C7BzO. Protein concentrations were determined by Bradford assay. Up to 25 µg of protein per sample were separated by SDS–PAGE, excised and subjected to in-gel tryptic digestion. Peptides were analysed by reverse-phase liquid chromatography coupled to high-resolution tandem mass spectrometry on an Orbitrap-based platform at the PISSARO proteomics facility, University of Rouen Normandy. Protein identification used Proteome Discoverer with MASCOT against the human Swiss-Prot database.

Proteins supported by at least two peptides were retained. Technical LC–MS/MS replicate measurements were evaluated for consistency. Feature-level processing and the initial statistical output, from which the 378 proteins were selected, were obtained in Progenesis, which could not average technical replicates; the Close–Far comparison was therefore repeated in Perseus at the level of independent patient samples after averaging technical replicates. Exploratory criteria were nominal p < 0.05 and |log2FC| ≥ 1 where indicated. IPA was used to prioritise functional and pathway hypotheses from differentially abundant proteins. Given the small sample size, pathway Z-scores and nominal protein-level p-values were not treated as confirmatory evidence.

### Multiplex immunofluorescence and spatial marker profiling

Twelve FFPE tumour sections (four per ventricular-distance group) were analysed using the Cell DIVE multiplex immunofluorescence workflow. After deparaffinisation, rehydration and antigen retrieval, sequential antibody staining and image acquisition were followed by dye inactivation, image registration to DAPI nuclei and correction for illumination and tissue autofluorescence. The broader panel included tumour/glial (SOX2, SOX9, GFAP), proliferation (Ki67), myeloid (CD163, CD68), lymphoid/regulatory (CD4, CD8, FOXP3, CD45RO, PD-1, PD-L1), and stromal/vascular (SMA/α-SMA, COL1A1) markers, alongside neuronal markers. Quantitative fractions were available for SOX2, CD163, CD68, SMA, COL1A1, Ki67, CD4 and CD8a, plus SOX2-positive proliferative and non-proliferative categories. Cell-level fractions were summarised within each biological specimen before calculating group means and standard deviations; individual cells and imaging fields were not treated as biological replicates. Antibody clones, dilutions and marker-positivity thresholds are available from the corresponding author upon request.

### Statistical analyses

Analyses were performed in R unless otherwise stated. Overall survival was compared using Kaplan–Meier estimates and log-rank tests. A univariable Cox proportional-hazards model evaluated tumour–ventricle distance as a continuous variable, and hazard ratios are reported per 5-mm increase with 95% confidence intervals. The Cox model included the 52 patients with complete data for the variables analysed. Kruskal–Wallis tests were used for non-parametric comparisons between groups, and multiple-testing correction used the Benjamini–Hochberg procedure where specified. Proteomic thresholds were nominal unless otherwise stated. Multiplex immunofluorescence results were summarised descriptively at specimen level (n = 4 per distance category), and no inferential tests were applied. No adjusted Cox model is presented; distance should therefore not be interpreted as an independent prognostic factor.

## DATA AVAILABILITY

Data supporting the reported findings include clinical variables, RNA-seq and WGS data, proteomic abundance matrices and Cell DIVE imaging-derived quantification. Public deposition or controlled-access arrangements for human-derived data will be described here after confirmation of consent, ethics and repository permissions.

## ACKNOWLEDGEMENTS

We are deeply grateful to the patients with glioblastoma and their families, whose generous participation made this study possible. We thank the Biobank of the General University Hospital of Castellón for providing samples and facilitating this work, and the Departments of Neurosurgery, Pathology, Radiology of the General University Hospital of Castellón as well as the Departments of Medical Oncology and Radiation Oncology of the Provincial Hospital Consortium of Castellón (Consorcio Hospitalario Provincial de Castellón), for their support with patient care, tissue collection, imaging and clinical follow-up data. We thank the Centro Nacional de Análisis Genómico (CNAG, Barcelona) for RNA and whole-genome sequencing services, and the PIS-SARO proteomics platform (Université de Rouen Normandie) for mass spectrometry analyses. The authors also acknowledge the use of the services and facilities of Universitat Jaume I (Castellón).

This study was funded by MCIN/AEI/10.13039/501100011033 and the European Union NextGenerationEU/PRTR (grant CPP2021-8817); by a UJI–FISABIO Preparatory Actions grant for Future Joint Research/Innovation Projects 2023 (“Cazafantasmas-glioblastoma: buscando exterminar la reaparición del glioblastoma después de los tratamientos convencionales; by Universitat Jaume I research project UJI-2023-29 (23I469; “The force awakens in GBM: molecular mechanisms that regulate quiescent cells in glioblastoma”); and by the European Digital UniverCity (EDUC 2025; “Unraveling Glioblastoma: Molecular Study of Quiescent Cells”).

J.L.G.-A. acknowledges an “EPRIEX” contract from the Spanish Recovery, Transformation and Resilience Plan and a UJI industrial doctorate fellowship in collaboration with FISABIO and cofunded by the Generalitat Valenciana and the European Social Fund, under the collaboration agreement between the Generalitat Valenciana (Department of Education, Culture, Universities and Employment) and Universitat Jaume I for the promotion of industrial doctorates. J.L.G.A acknowledges an-Erasmus+ Traineeship grant (Erasmus+ Programme). P.G.-P. received a predoctoral fellowship from Universitat Jaume I (grant 19I001.03 UJI-2025-28). E.C.-M. acknowledges a fellowship from the Spanish Association Against Cancer (AECC). M.A.M.-T. was supported by a “María Zambrano” research contract (MAZ/2021/03 UP2021-021) funded by the European Union–NextGenerationEU and by a Ramón y Cajal contract (RYC2022-038481-I) funded by MCIN/AEI/10.13039/501100011033. The authors declare no competing interests. The funders had no role in the design of the study; in the collection, analysis or interpretation of the data; in the writing of the manuscript; or in the decision to publish the results.

## Author contributions

Conceptualization: J.L.G.-A., M.A.M.-T.

Methodology: J.L.G.-A., M.A.M.-T.

Formal analysis: J.L.G.-A.

Bioinformatic analysis (RNA-seq, WGS and integration): J.L.G.-A., C.C., S.R.

Proteomics: J.L.G.-A., C.G., P.C.

Multiplex immunofluorescence and image analysis: J.L.G.-A., J.B., O.M.

Investigation: J.L.G.-A., P.G.-P., E.C.-M., M.B.F., L.G.G.-B., C.M.-C., M.A.M.-T.

Resources (patient samples and clinical data): J.L.G.-A., L.G.G.-B.

Data curation: J.L.G.-A.

Visualization: J.L.G.-A.

Writing – original draft: J.L.G.-A.

Writing – review & editing: all authors

Supervision: C.M.-C., M.A.M.-T.

Funding acquisition: M.A.M.-T.

Project administration: M.A.M.-T.

All authors read and approved the final manuscript.

